# Haplotype-resolved chromosome-level genome assembly of four European white oak species

**DOI:** 10.64898/2026.08.20.745905

**Authors:** Gabriele Magris, Camilla Avanzi, Francesca Bagnoli, Ludovic Duvaux, Elodie Belmonte, Giovanni Giuseppe Vendramin, Andrea Piotti, Sara Pinosio

**Author notes:** Corresponding author: Sara Pinosio.

## Abstract

European white oaks (*Quercus* section *Quercus*) are ecologically and economically important forest trees characterized by extensive shared genetic variation and a history of interspecific gene flow. Genomic resources remain uneven across species, limiting comparative analyses and pangenome development. Here, we present haplotype-resolved chromosome-scale genome assemblies and genome annotations for four European white oak species: *Quercus robur, Q. petraea, Q. pubescens*, and *Q. frainetto*. The assemblies were generated from PacBio HiFi sequencing data and include both phased haplotypes for each species. Genome sizes range from 779 to 817 Mb and all assemblies are organized into 12 chromosome-scale pseudomolecules with high completeness and contiguity. We additionally provide species-specific repeat annotations, structurally and functionally annotated protein-coding gene sets, and complete organellar genomes. The dataset includes the first reference genomes for *Q. pubescens* and *Q. frainetto*, together with newly generated assemblies for *Q. robur* and *Q. petraea* produced using a consistent sequencing and analysis workflow. These resources provide a standardized framework for comparative genomics, pangenome construction, genome evolution studies, and investigations of adaptation and introgression across European white oaks.

## Background & Summary

Oaks are key forest tree species broadly distributed throughout the Northern Hemisphere, where they represent valuable natural resources from economic, ecological, social, and historical perspectives. Processes such as hybridization and adaptive introgression have been identified as fundamental factors underlying their evolutionary success^1^ so much so that the concept of syngameon was introduced to describe “the persistence of interfertile species that remain largely distinct despite living in sympatry and having numerous opportunities for hybridization”^2^. This evolutionary dynamic is particularly evident within the white oaks (section *Quercus*, subgenus *Quercus*), one of the largest and most ecologically successful lineages of the genus.

White oaks are foundation species of several forest ecosystems in Europe, ranging from endangered lowland forests threatened by urbanization to forests typical of Mediterranean environments. *Quercus robur, Q. petraea, Q. pubescens* and *Q. frainetto*, are among the most relevant and widespread members of the Mediterranean white oaks syngameon, ordered here from the most drought-sensitive to the most drought resistant^3^. The persistence of such a group of otherwise distinct species interconnected by limited gene flow is stimulating research on oak pan-genomes. In fact, whole-genome sequencing has shown that inter-specific gene flow can occur without disrupting species integrity, which is restricted to a limited part of the genome, maintaining species barriers in European white oaks^1^.

The availability of chromosome-scale reference genomes has greatly advanced studies on oak evolution, adaptation, and structural genomic variation. However, genomic resources remain unevenly distributed across European white oaks, limiting comparative analyses of structural variation, haplotype diversity, and the genomic consequences of historical and ongoing introgression. Moreover, the limited taxonomic representation and methodological heterogeneity of currently available assemblies still hamper efforts to capture the full extent of genomic diversity within the European white oak syngameon. Expanding the number of species represented by high-quality, haplotype-resolved assemblies is therefore a key prerequisite for building a comprehensive pangenome of European white oaks. This need is particularly relevant for Mediterranean populations, which harbor a substantial fraction of the evolutionary history and genetic diversity of European oaks. Southern European refugia acted as long-term reservoirs of diversity during Quaternary climatic oscillations and remain hotspots of intraspecific variation and adaptive potential^4–6^. Comprehensive genomic resources from representative Mediterranean taxa are therefore essential for investigating the origin, maintenance, and adaptive significance of genomic diversity across the European white oak syngameon.

To expand the genomic representation of European white oaks and provide a consistent resource for comparative analyses, we generated high-quality haplotype-resolved de novo genome assemblies and structural annotations for four European white oak species (*Quercus pubescens, Q. robur, Q. petraea*, and *Q. frainetto*). The assemblies of *Q. pubescens* and *Q. frainetto* provide the first reference genomes for these species, while those of *Q. robur* and *Q. petraea* complement existing genomic resources with new assemblies generated using the same sequencing and analytical workflow across all four species. This standardized approach minimizes technical biases and provides a robust framework for comparative genomics, pangenome construction, and investigations of hybridization, adaptation, and genome evolution in European white oaks.

## Methods

### Sampling, Library Preparation, and Sequencing

Terminal branches (∼15 cm in length) were collected from naturally occurring adult trees of *Quercus frainetto, Q. petraea, Q. pubescens* and *Q. robur* in southern Italy. Sampling sites were located at Rotondella (40.1684° N, 16.5580° E) for *Q. pubescens*, Bosco Pantano (40.1682° N, 16.6875° E) for *Q. robur*, Noepoli (40.0656° N, 16.3140° E) for *Q. frainetto*, and Rifreddo (40.5563° N, 15.8220° E) for *Q. petraea*. DNA extraction and single-molecule real-time long-read sequencing were performed at the Gentyane Sequencing Platform (Clermont-Ferrand, France). DNA was extracted from leaf buds (1 g) using the PacBio® Nanobind® HMW DNA Extraction Kit, following the protocols “Isolating nuclei from plant tissue using TissueRuptor disruption” and “Extracting HMW DNA from plant nuclei using Nanobind® kits”. High-quality PacBio HiFi (CCS) reads were generated with a PacBio Revio System Sequencer (Pacific Biosciences, Menlo Park, CA, USA). On average, 5.8 M reads were generated per sample, corresponding to a total of 99.1 M base pairs and a depth of sequencing ranging from 110 to 130X (considering a genome length of 820Mb). N50 of reads ranged from 15.4 to 16.9 kb.

### Genome assembly and construction of chromosome pseudomolecules

High-quality chromosome-scale genome assemblies were generated for both phased haplotypes of *Quercus robur, Q. pubescens, Q. petraea*, and *Q. frainetto* using hifiasm^7^ v0.25.0-r726 (Table 1). The contig assemblies ranged from 787.0 Mb (*Q. robur* Hap2) to 859.1 Mb (*Q. petraea* Hap1). Assembly contiguity was consistently high across all haplotypes and species with contig N50 values ranging from 37.5 Mb in *Q. pubescens* Hap2 to 58.2 Mb in *Q. pubescens* Hap1. The largest contigs reached 104.7 Mb in *Q. robur* Hap1. Although the number of contigs varied substantially among assemblies, from 130 in *Q. robur* Hap2 to 646 in *Q. petraea* Hap1, all genomes exhibited high continuity, demonstrating the ability of long-read sequencing to reconstruct highly contiguous phased haplotypes.

**Table 1.** Summary statistics of chromosome-scale phased *Quercus* genome assemblies.

| Metric | *Q. frainetto* |  | *Q. petraea* |  | *Q. pubescens* |  | *Q. robur* |  |
| --- | --- | --- | --- | --- | --- | --- | --- | --- |
|  | Hap1 | Hap2 | Hap1 | Hap2 | Hap1 | Hap1 | Hap2 | Hap1 |
| **ASSEMBLY** |  |  |  |  |  |  |  |  |
| Contig assembly size (Mb) | 839.56 | 805.74 | 859.13 | 809.77 | 831.69 | 831.2 | 823.05 | 787.01 |
| Chromosome assembly size (Mb) | 816.92 | 796.23 | 804.22 | 799.88 | 798.6 | 804.09 | 799.87 | 779.01 |
| Anchored bases (%) | 97.3 | 98.82 | 93.61 | 98.78 | 96.98 | 96.74 | 97.18 | 98.98 |
| GC (%) | 35.85 | 35.71 | 35.92 | 35.74 | 35.86 | 35.7 | 35.77 | 35.63 |
| **ASSEMBLY CONTIGUITY** |  |  |  |  |  |  |  |  |
| Number of contigs | 395 | 142 | 646 | 134 | 559 | 168 | 391 | 130 |
| Contig N50 (Mb) | 54.94 | 50.4 | 55.86 | 50.76 | 58.18 | 37.5 | 54.99 | 43.44 |
| Largest contig (Mb) | 95.03 | 88.11 | 89.87 | 102.64 | 103.15 | 72.1 | 104.69 | 67.35 |
| Chromosomes | 12 | 12 | 12 | 12 | 12 | 12 | 12 | 12 |
| Scaffold N50 (Mb) | 69.24 | 69.97 | 68.97 | 66.39 | 67.59 | 67.07 | 69.48 | 67.35 |
| **ASSEMBLY QUALITY** |  |  |  |  |  |  |  |  |
| LAI | 30.59 | 30.67 | 33.36 | 32.02 | 32.08 | 31.58 | 28.56 | 28.52 |

The contigs of the eight haplotypes were subsequently anchored, ordered and oriented using the *Q. robur* dhQueRobu3.1 assembly (hereafter Qrob3.1, GCA_932294415.1) as the reference with RagTag v2.1.0^8^. RagTag was used to anchor, order and orient the hifiasm-generated contigs, without modifying their underlying structure (i.e. without splitting, merging, or otherwise altering the contigs). Thus, the haplotype-specific contigs generated by hifiasm were retained as independent units, and the reference-guided step was used only to establish their chromosome-scale order and orientation. Qrob3.1 provides greater sequence completeness and contiguity than the original *Q. robur* reference genome^9^, although the orientation of a few chromosomes differs between the two assemblies (Supplementary Figure 1A). A chromosome-scale assembly of *Q. petraea* has also been recently released; however, its chromosomes are ordered by size rather than according to the Qrob3.1 chromosome nomenclature (Supplementary Figure 1B).

Following chromosome anchoring to Qrob3.1 reference genome, the final chromosome-scale assemblies ranged between 779.0 Mb and 816.9 Mb, with all genomes assembled into the expected 12 pseudomolecules corresponding to the haploid chromosome number of white oaks. Scaffold N50 values ranged from 66.4 Mb (*Q. petraea* Hap2) to 70.0 Mb (*Q. frainetto* Hap2), reflecting near chromosome-arm continuity across all assemblies. Genome GC content was remarkably conserved, varying only between 35.6% and 35.9%, consistent with previous genome assemblies reported for the genus *Quercus*^10,11^. Complete organelle genomes were reconstructed using Oatk^12^ and incorporated into the first haplotype assembly of each species.

### Repetitive elements annotation

Repetitive sequences were annotated in the assembled genomes using EDTA^13^, revealing highly similar repeat landscapes across the eight phased haplotypes (Table 2). Tandem repeats were identified separately using a modified version of Tandem Repeats Finder (https://github.com/lh3/TRF-mod). Repetitive DNA accounted for 52.7% to 54.2% of the assembled genomes, with only minor differences between haplotypes within each species. These values are consistent with those reported for other chromosome-scale *Quercus* genomes, including *Q. lobata* (54%), *Q. mongolica* (53.75%), *Q. rubra* (47.51%), *Q. gilva* (54.17%), *Q. glauca* (59%), *Q. canariensi*s (54.35%) and *Q. alba* (58% and 59% for HapA and HapB, respectively)^10,14–19^.

**Table 2.** Summary of major transposable element (TE) superfamilies in phased *Quercus* genome assemblies.

| Metric | *Q. frainetto* |  | *Q. petraea* |  | *Q. pubescens* |  | *Q. robur* |  |
| --- | --- | --- | --- | --- | --- | --- | --- | --- |
|  | Hap1 | Hap2 | Hap1 | Hap2 | Hap1 | Hap2 | Hap1 | Hap2 |
| Assembly size (Mb) | 816.9 | 796.2 | 804.2 | 799.9 | 798.6 | 804.1 | 799.9 | 779.0 |
| Genome masked (%) | 54.23 | 54.15 | 52.73 | 52.95 | 54.16 | 53.29 | 53.51 | 53.42 |
| L1 (%) | 2.37 | 2.08 | 2.02 | 2.17 | 1.86 | 2.08 | 2.15 | 2.17 |
| Copia (%) | 6.77 | 6.77 | 5.59 | 5.31 | 5.16 | 6.41 | 6.54 | 6.70 |
| Gypsy (%) | 7.91 | 9.07 | 8.61 | 8.54 | 7.89 | 8.18 | 12.00 | 11.50 |
| Unknown LTR (%) | 14.36 | 13.25 | 13.23 | 14.77 | 14.85 | 13.90 | 13.89 | 13.33 |
| CACTA (%) | 0.94 | 0.82 | 0.98 | 1.16 | 0.91 | 0.70 | 0.70 | 1.02 |
| Mutator (%) | 3.46 | 3.62 | 3.57 | 2.82 | 3.17 | 7.11 | 3.43 | 3.19 |
| PIF/Harbinger (%) | 2.66 | 2.30 | 2.54 | 2.49 | 2.24 | 2.33 | 2.23 | 2.61 |
| hAT (%) | 2.47 | 2.50 | 2.44 | 2.97 | 2.34 | 2.25 | 2.69 | 2.56 |

Long terminal repeat (LTR) transposons were the dominant class of repetitive elements in all assemblies, with Gypsy, Copia and unknown LTR elements accounting for the largest fraction of the genomes, in agreement with previous studies. Among DNA transposons, Mutator elements were the most abundant superfamily occupying approximately 2.8-3.6% of the genome in most haplotypes. An exception was *Q. pubescens* Hap2, where Mutator sequences accounted for 7.1% of the genome.

### Gene prediction and functional annotation

Protein-coding genes were identified and annotated in each phased haplotype by integrating *ab initio* gene prediction, protein homology, and transcriptome evidence. Prior to annotation, genome assemblies were soft masked using the repeat libraries generated with EDTA. Gene prediction was performed with BRAKER3^20^, integrating RNA-seq evidence to improve the accuracy of gene models. Publicly available Illumina RNA-seq reads (Supplementary Table 1) were aligned to each haplotype using HISAT2^21^ and the resulting alignments were assembled into transcript models with StringTie^22^.

Gene models were subsequently refined using Evian^23^, which integrates transcript evidence to improve structural annotation. Publicly available Illumina RNA-seq data were combined with full-length Oxford Nanopore Technologies (ONT) RNA sequencing reads to define 5’ and 3’ untranslated regions (UTRs), resulting in more complete gene models and more accurate annotation. To remove likely TE-derived predictions, the final gene set was filtered using the species-specific TE annotation. Gene models whose coding sequences (CDS) overlapped annotated transposable elements by ≥80% of their total CDS length were discarded.

The resulting gene annotations were highly consistent across the four *Quercus* species, indicating a robust and uniform annotation (Table 3). The number of predicted protein-coding genes ranged from approximately 40,700 in *Q. petraea* to approximately 42,000 in *Q. robur* and *Q. frainetto* haplotypes. On average each gene produced 1.9 transcript isoforms, suggesting a moderate level of alternative splicing conserved among species. Gene architecture was similarly conserved across the dataset. Mean gene length ranged between 6.3 and 6.7 kb, with the average coding sequence length close to 1.2 kb in every haplotype. Predicted transcripts contained on average 5.5-5.6 exons and between 43,677 and 44,631 transcripts showed both 5’ and 3’ UTR annotations. Given the modest differences in genome size and repeat content, the four oak genomes displayed remarkably similar gene content and structural organization, supporting the completeness and consistency of the annotation pipeline. Functional annotation of the predicted proteins was performed using OmicsBox v4.0.54 (BioBam, Valencia, Spain). Predicted protein sequences were searched against the NCBI non-redundant protein database using DIAMOND BLAST, while conserved protein domains and signatures were identified with InterProScan. Gene Ontology (GO) annotations were assigned through the GO Mapping and GO Annotation pipelines implemented in OmicsBox, combining homology-based and domain-based evidence. Enzyme Commission (EC) identifiers were assigned using the EC Mapping workflow.

**Table 3.** Summary of gene prediction in phased *Quercus* genome assemblies.

| Metric | *Q. frainetto* |  | *Q. petraea* |  | *Q. pubescens* |  | *Q. robur* |  |
| --- | --- | --- | --- | --- | --- | --- | --- | --- |
|  | Hap1 | Hap2 | Hap1 | Hap2 | Hap1 | Hap2 | Hap1 | Hap2 |
| **Gene content** |  |  |  |  |  |  |  |  |
| Genes | 41,805 | 41,584 | 41,418 | 40,715 | 41,572 | 41,320 | 41,810 | 40,907 |
| Transcripts | 78,265 | 77,640 | 77,365 | 76,930 | 77,101 | 77,555 | 78,137 | 76,525 |
| Mean transcripts per gene | 1.9 | 1.9 | 1.9 | 1.9 | 1.8 | 1.9 | 1.9 | 1.9 |
| Single-exon genes | 13,453 | 13,673 | 13,417 | 12,701 | 13,670 | 13,444 | 13,711 | 13,010 |
| **UTR annotation** |  |  |  |  |  |  |  |  |
| Transcripts with both UTRs | 44,539 | 44,430 | 44,068 | 44,580 | 43,677 | 44,538 | 44,631 | 44,085 |
| Transcripts with at least one UTR | 46,849 | 46,641 | 46,337 | 46,846 | 45,874 | 46,760 | 46,821 | 46,273 |
| **Gene structure** |  |  |  |  |  |  |  |  |
| Mean gene length (kb) | 6.75 | 6.39 | 6.59 | 6.75 | 6.33 | 6.71 | 6.45 | 6.48 |
| Mean transcript length (kb) | 7.87 | 7.72 | 7.63 | 7.83 | 7.54 | 7.86 | 7.73 | 7.47 |
| Mean CDS length (kb) | 1.20 | 1.20 | 1.21 | 1.21 | 1.20 | 1.20 | 1.20 | 1.21 |
| Mean exons per transcript | 5.6 | 5.6 | 5.5 | 5.6 | 5.5 | 5.6 | 5.6 | 5.6 |

Genome-wide patterns of repetitive sequences and gene density were represented using Circos v0.69-8^24^. Comparable patterns were observed across all four *Quercus* species (Figure 1), where repetitive sequences were enriched in pericentromeric regions, while gene density was higher along chromosome arms. Tandem repeat analysis identified a highly abundant 146 bp repeat within centromeric regions. This repeat was then used as custom RepeatMasker library (http://www.repeatmasker.org) to annotate and mask centromeric regions across the genomes.

**Figure 1.**
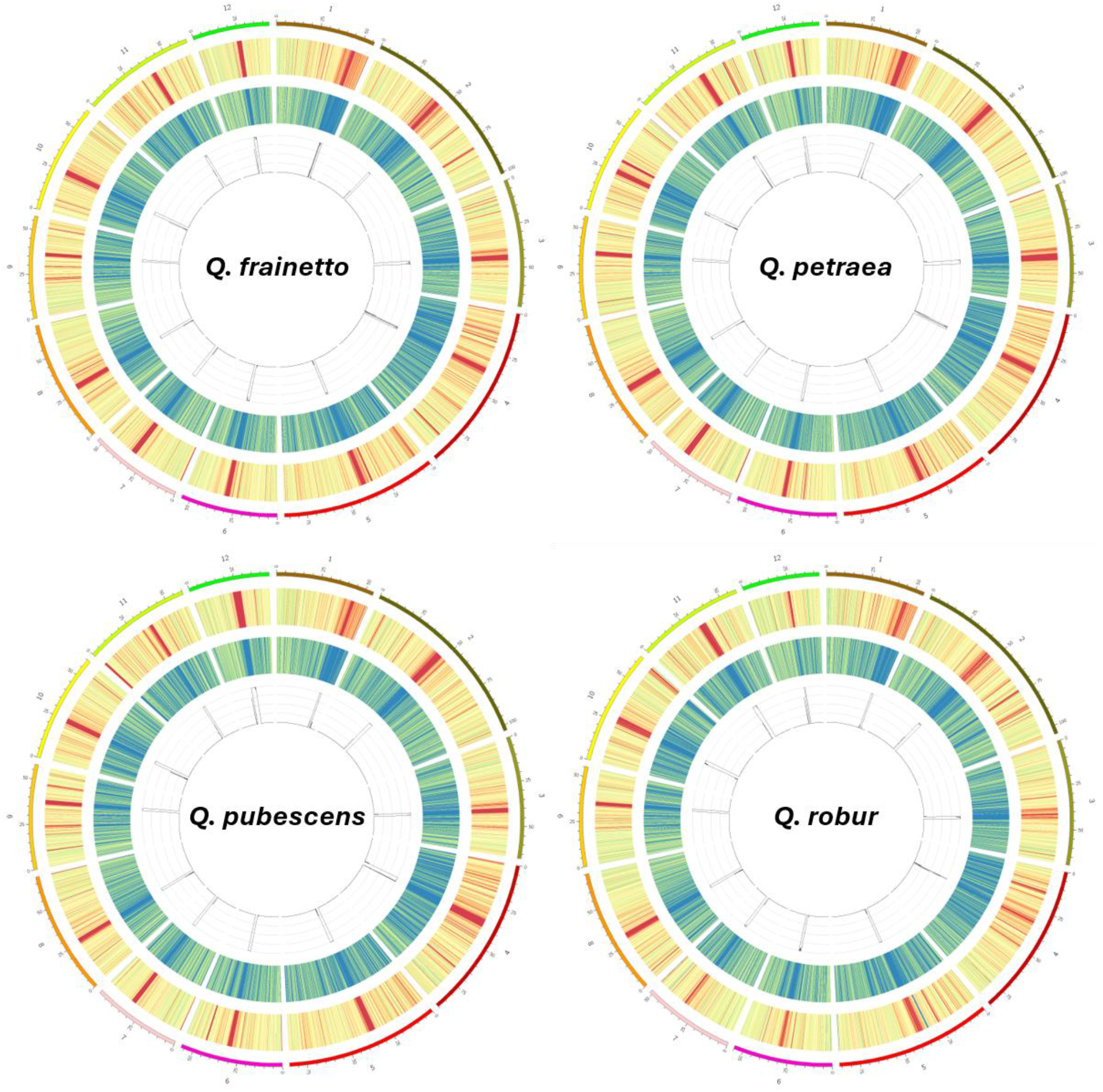
Chromosome-wide distribution of repetitive, coding, and centromeric sequences in haplotype 1 assemblies of *Quercus*. Circos plots showing the genomic organization of *Q. frainetto, Q. petraea, Q. pubescens* and *Q. robur* (top-left to bottom-right). Chromosomes are represented by the outer colored ideograms and numbered according to the reference genome assembly. The outer heatmap represents repeat density, with the color scale ranging from yellow (lower density) to red (higher density), whereas the inner heatmap represents gene density, with the color scale ranging from blue (lower density) to yellow (higher density). The innermost histogram represents the abundance of centromeric repeat sequences. All features were calculated in non-overlapping 100 Kb genomic windows.

## Data Records

Raw sequencing data generated in this study, including PacBio HiFi and ONT RNA-seq reads, have been deposited in the NCBI Sequence Read Archive (SRA). ONT-RNA-seq data were deposited under the BioProject PRJNA1496551^25^, with the accession numbers SRR39678451, SRR39678452, SRR39678453, SRR39678454, for *Q. petraea, Q. pubescens, Q. robur* and *Q. frainetto*, respectively. PacBio Hifi reads were deposited under the BioProject PRJNA1498310^26^. The final haplotype-resolved genome assemblies have been deposited in the NCBI genome database. *Quercus frainetto* haplotypes 1 and 2 were deposited under BioProjects PRJNA1489469^27^ (JCAOMP000000000) and PRJNA1489468^28^ (JCAOMQ000000000), respectively, with locus tag prefixes (LTPs) AC5KOV and AC5KOU. *Quercus petraea* haplotypes 1 and 2 were deposited under PRJNA1489465^29^ (JCAOML000000000) and PRJNA1489464^30^ (JCAOMM000000000) with LTPs AC5KOR and AC5KOQ, *Quercus pubescens* haplotypes 1 and 2 under PRJNA1489467^31^ (JCAOMN000000000) and PRJNA1489466^32^ (JCAOMO000000000) with LTPs AC5KOT and AC5KOS, and *Quercus robur* haplotypes 1 and 2 under PRJNA1489463^33^ (JCAOMJ000000000) and PRJNA1489462^34^ (JCAOMK000000000) with LTPs AC5KOP and AC5KOO, respectively. Genome annotation files have been deposited in figshare (10.6084/m9.figshare.33122921).

## Technical Validation

Genome completeness was evaluated using BUSCO v6.0.0^35^ with the embryophyta dataset, complemented by k‐mer based analyses performed with KAT‐comp (k = 31). All four *de novo* genome assemblies showed consistently high completeness (≥99.4% complete BUSCOs), representing a clear improvement over the earlier *Q. robur* PM1N reference genome (95.5% completeness; Plomion et al., 2018) and comparable to recent high-quality oak assemblies generated by the Darwin Tree of Life Project (*Quercus petraea*, GCA_964102825.1; *Quercus robur*, GCA_932294415.1) (Figure 2). Most BUSCO genes were recovered as complete and single-copy (∼94.7–95.1%), with only a small proportion of duplicated genes (∼4.6–5.1%) and negligible fragmented or missing components.

**Figure 2.**
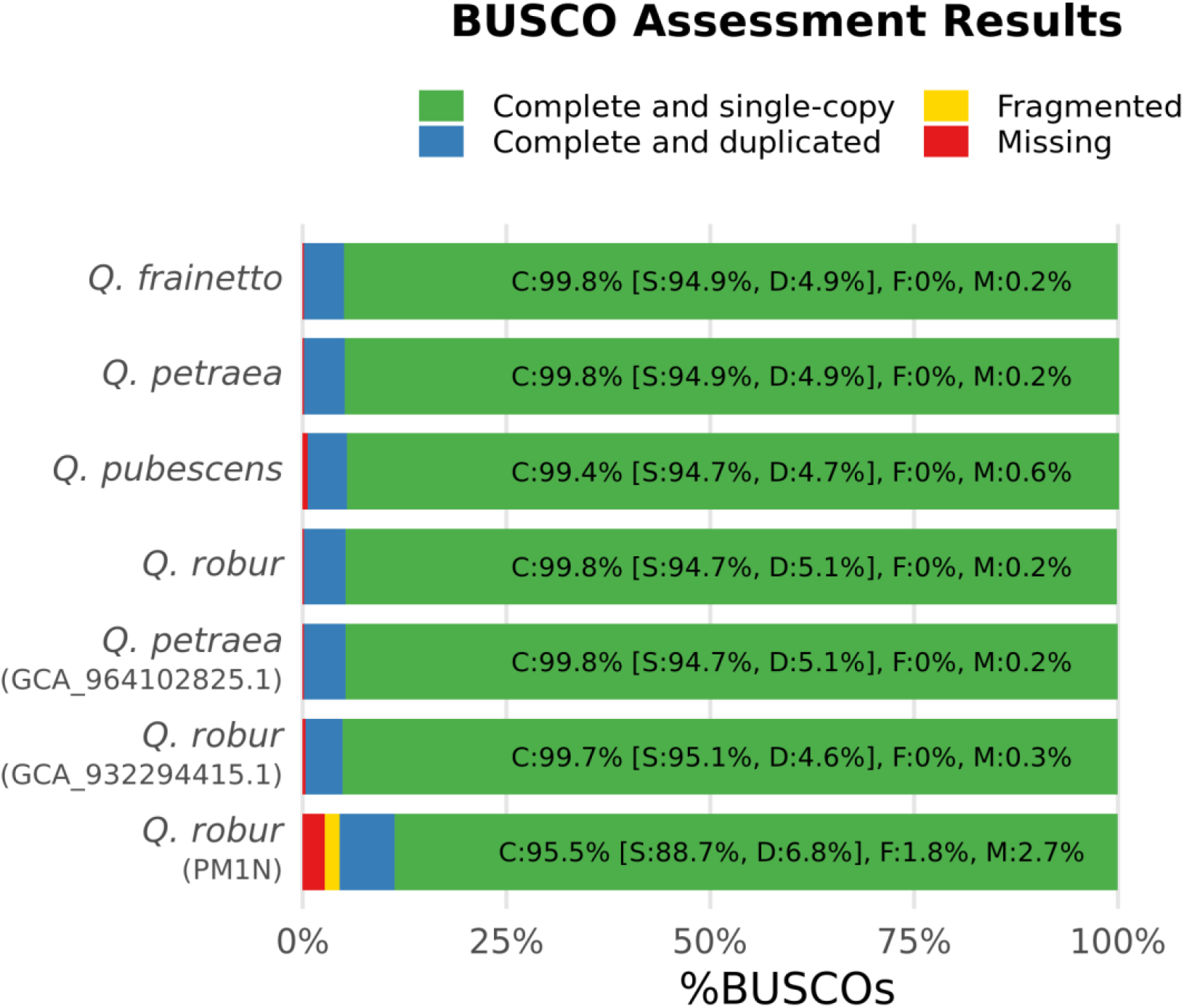
BUSCO completeness assessment of *Quercus* genome assemblies. BUSCO analysis performed using the embryophyta dataset on the four newly assembled genomes (*Q. frainetto, Q. petraea, Q. pubescens*, and *Q. robur*) and compared with previously published references. Bars represent the proportions of complete single-copy (green), complete duplicated (blue), fragmented (yellow), and missing (red) BUSCO genes.

K‐mer spectrum comparisons between HiFi sequencing reads and the haplotype-resolved assemblies further support these results (Figure 3). Analyses were conducted on the primary haplotypes (Hap1), with secondary haplotypes (Hap2) showing comparable profiles (data not shown). All Hap1 assemblies exhibit the expected bimodal k‐mer distribution reflecting heterozygous and homozygous sequence content, with the majority of k‐mers recovered as single-copy (1x k‐mers), indicating accurate representation of the sequencing data. Missing k‐mers (0x k‐mers) are limited and largely restricted to low multiplicity values, consistent with sequencing noise, while the low proportion of duplicated k‐mers (2x k‐mers) indicates minimal redundancy and limited assembly artefacts.

**Figure 3.**
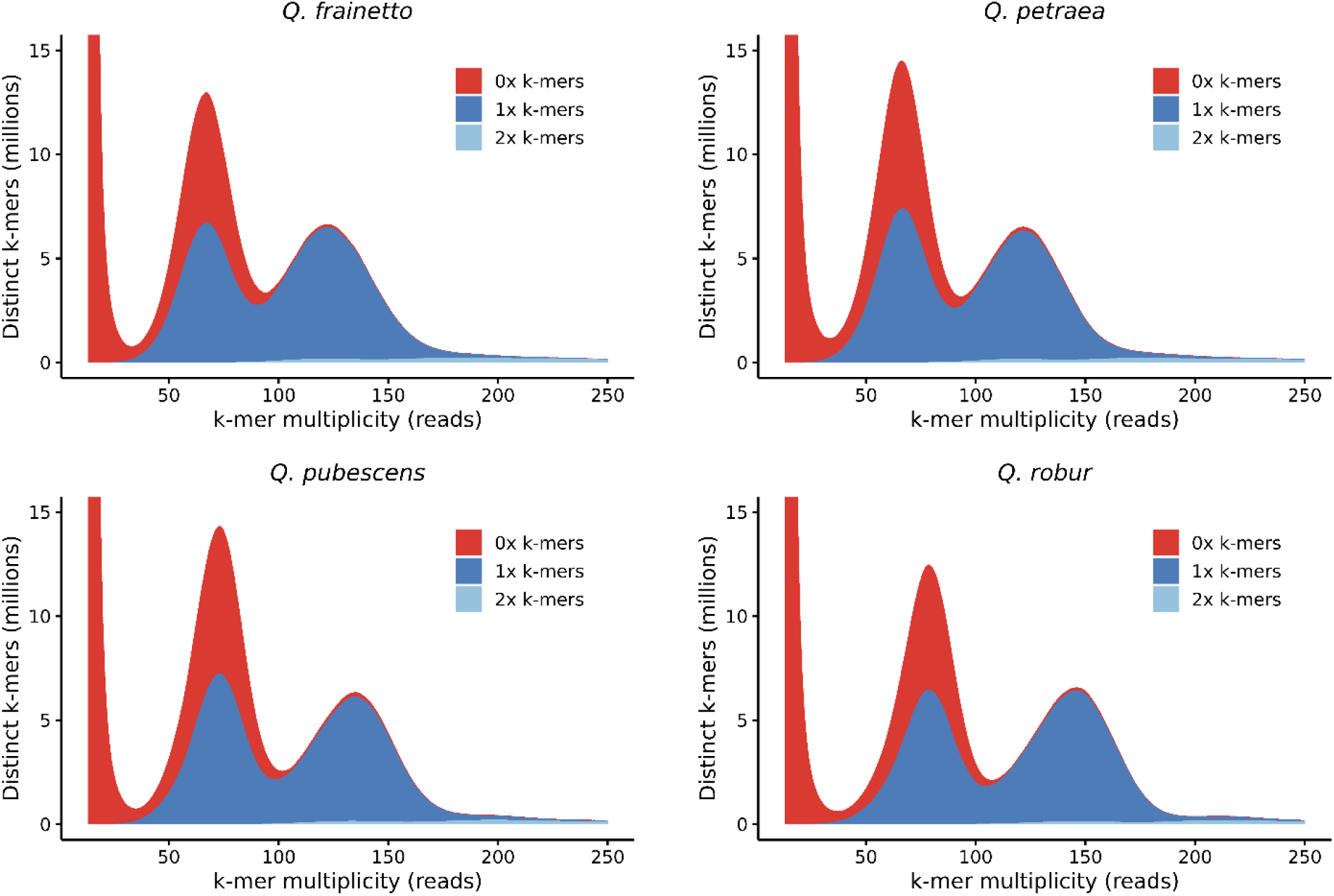
K‐mer spectrum comparison between HiFi reads and haplotype‐resolved genome assemblies. KAT‐comp analyses (k = 31) comparing HiFi sequencing reads to the primary haplotype assemblies (Hap1) of *Q. frainetto, Q. petraea, Q. pubescens*, and *Q. robur*.

Assembly quality within highly repetitive regions was assessed using the LTR Assembly Index (LAI)^36^. All assemblies largely exceeded the reference‐quality threshold (LAI ≥ 20), indicating an accurate reconstruction of repetitive sequences (Table 1). Telomeric repeat analysis performed with tidk^37^ further supported the high completeness of the assemblies, with 10 to 12 chromosomes in each haplotype displaying telomeric repeats at both ends. Comparative whole-genome synteny analyses between the two phased haplotypes of each species were obtained using SyRI v1.7.1^38^, and further reproduced graphically with plotsr v1.1.1^39^. High degree of chromosomal collinearity was observed across all species, with a limited number of inversions, duplications and translocations (Figure 4). Overall, the eight genomes represent highly complete chromosome-level assemblies with structural contiguity approaching telomere-to-telomere quality. Together, these results consistently indicate that the assemblies are highly complete and accurately capture the underlying sequence composition, supporting their reliability for downstream comparative and population genomic analyses.

**Figure 4.**
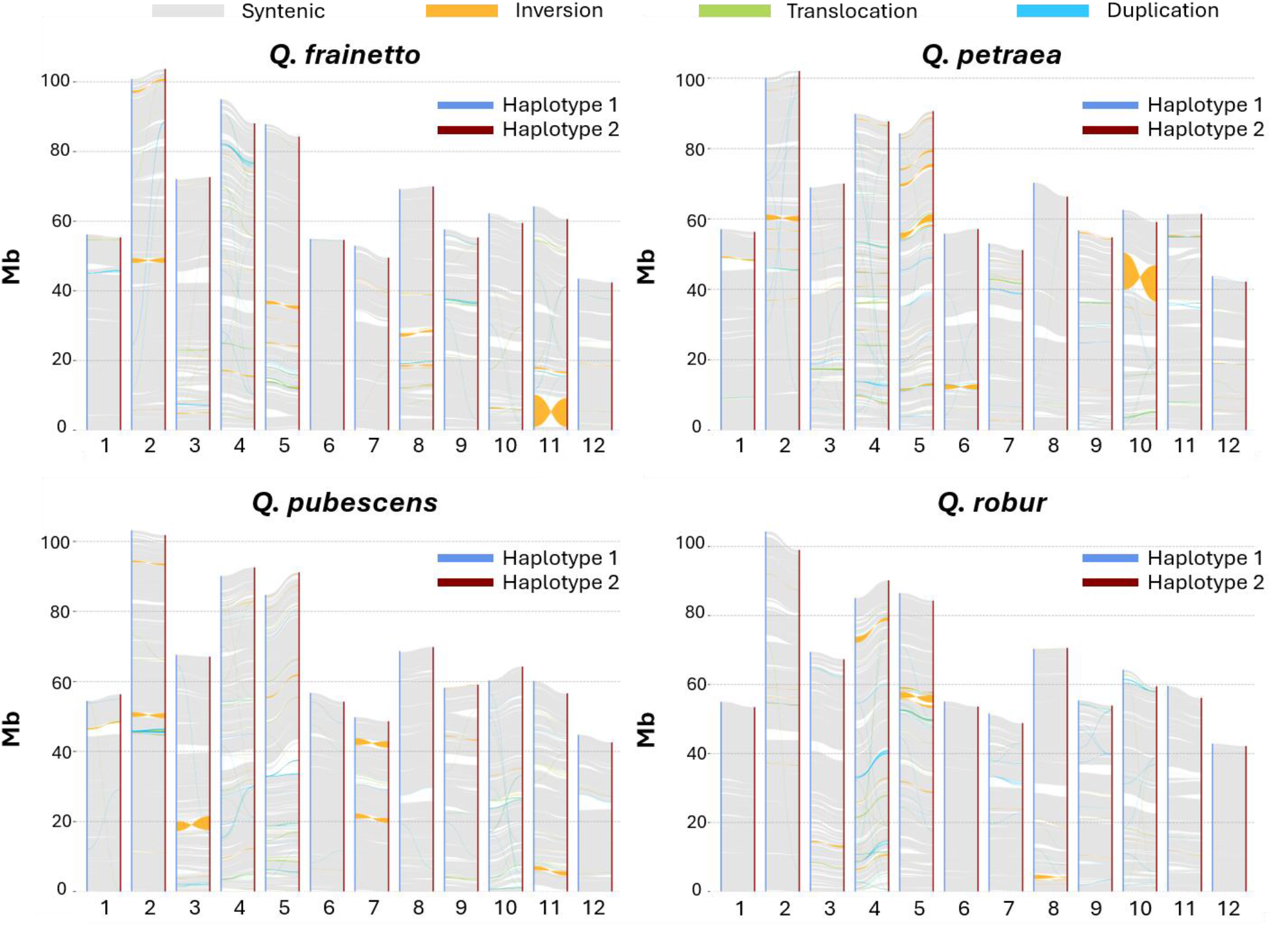
Structural synteny between haplotypes of *Quercus*. Genome-wide structural comparison between haplotype 1 and haplotype 2 of *Q. frainetto, Q. petraea, Q. pubescens and Q. robur* (top-left to bottom-right). Gray ribbons represent syntenic regions, orange ribbons represent inversions, green ribbons represent translocations and blue ribbons represent duplications.

## Data Availability

Raw sequencing data generated in this study have been deposited in the NCBI Sequence Read Archive (SRA) with accession numbers as follows. ONT-RNA-seq data were deposited under the BioProject PRJNA1496551. PacBio Hifi reads were deposited under the BioProject PRJNA1498310. Genome assemblies are accessible in NCBI under Bioproject PRJNA148946927 (*Q. frainetto* HAP1), PRJNA148946828 (*Q. frainetto* HAP2), PRJNA148946529 (*Q. petraea* HAP1), PRJNA148946430 (*Q. petraea* HAP2), PRJNA148946731 (*Q. pubescens* HAP1), PRJNA148946632 (Q. pubescens HAP2), PRJNA148946333 (*Q. robur* HAP1), PRJNA148946234 (*Q. robur* HAP2). The genome annotation files are available via Figshare at https://figshare.com/s/a575ab6287c89d1ab095.

## Code availability

No custom code was developed for this study. All bioinformatic analyses were performed using established software and tools following the corresponding documentation and recommended procedures. Software versions and the parameters adopted are reported in the Methods section.

## Acknowledgements

This research was funded by the Italian Ministry of University and Research (MUR) through the PRIN 2022 programme (*Progetti di Ricerca di Rilevante Interesse Nazionale*), project 2022BREB2H, entitled *“A pan-genomic approach to study local adaptation in Mediterranean oak forests (PanBiOak)”*. We acknowledge the GENTYANE platform (INRAE Clermont-Ferrand; DOI: 10.15454/1.5572409592543596E12) for NGS data production support. We thank Lucio Taverna and all the staff of the Friuli Venezia Giulia Regional Forest Nursery “Pascul” (https://www.regione.fvg.it/rafvg/cms/RAFVG/economia-imprese/agricoltura-foreste/foreste/FOGLIA10/) for providing the *Quercus petraea, Q. pubescens*, and *Q. robur* seedlings used in this study for ONT RNA sequencing, Catia Boggi and Maria Beatrice Castellani for providing technical assistance at various stages of the research, Maurizio Rosito and the staff of the Oasi WWF di Bosco Pantano for help in collecting *Q. robur* material, Antonio Lapolla, Francesco Ripullone and Michele Colangelo (UNIBAS) for indications about *Q. petraea* stands in the Basilicata region.

## Author contributions

S.P. and A.P. conceived the project and led the research. C.A., F.B., L.D., A.P., and G.G.V. were involved in sample collection and preparation. E.B. performed the extraction and sequencing experiments. G.M. and S.P. contributed to the bioinformatic analyses. G.M., S.P., and A.P. drafted the manuscript. All authors reviewed, revised, and approved the final manuscript.

