## Supplementary Figures and Tables for "Haplotype-resolved chromosome-level genome assembly of four European white oak species"

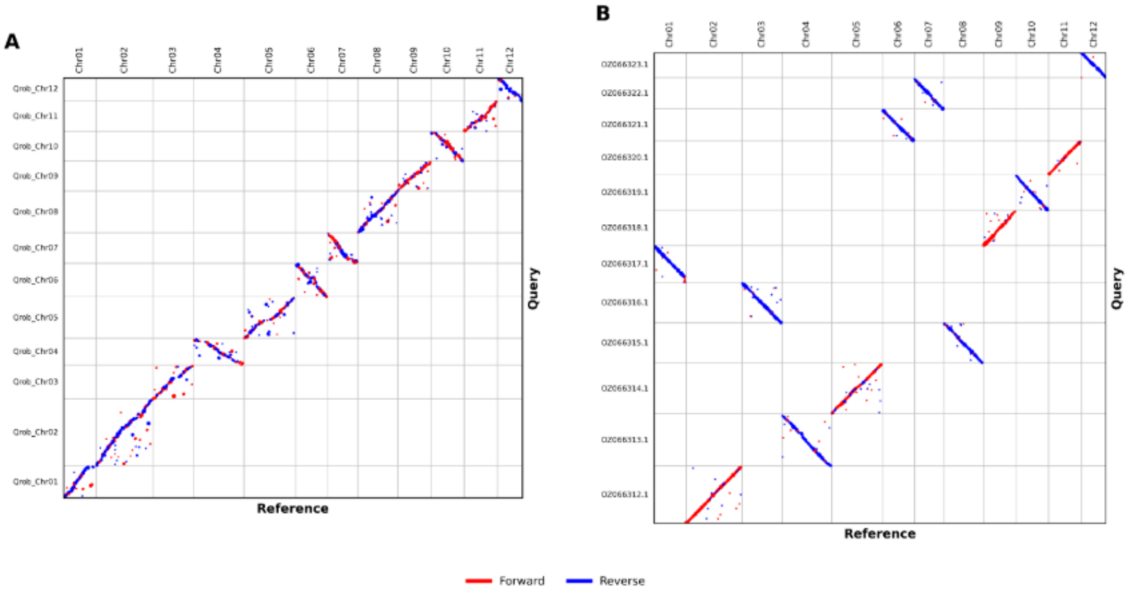


**Supplementary Figure 1. Whole genome comparisons between oak genome assemblies.** Whole genome pairwise alignments between the chromosome-scale assembly of *Quercus robur* (Qrob3.1, GCA_932294415.1), used as reference, and *Q. robur* PM1N (**A**) and *Q. petraea* (dhQuePetr1.hap1.1, GCA_964102825.1) (**B**) assemblies. Chromosomes along the x-axis correspond to the Qrob3.1 reference assembly. Only high-confidence alignments (≥10 kb and ≥95% nucleotide identity) are shown after retaining the best alignment per genomic bin and merging adjacent collinear alignment segments to improve readability. Forward alignments are shown in red, whereas reverse alignments are shown in blue.

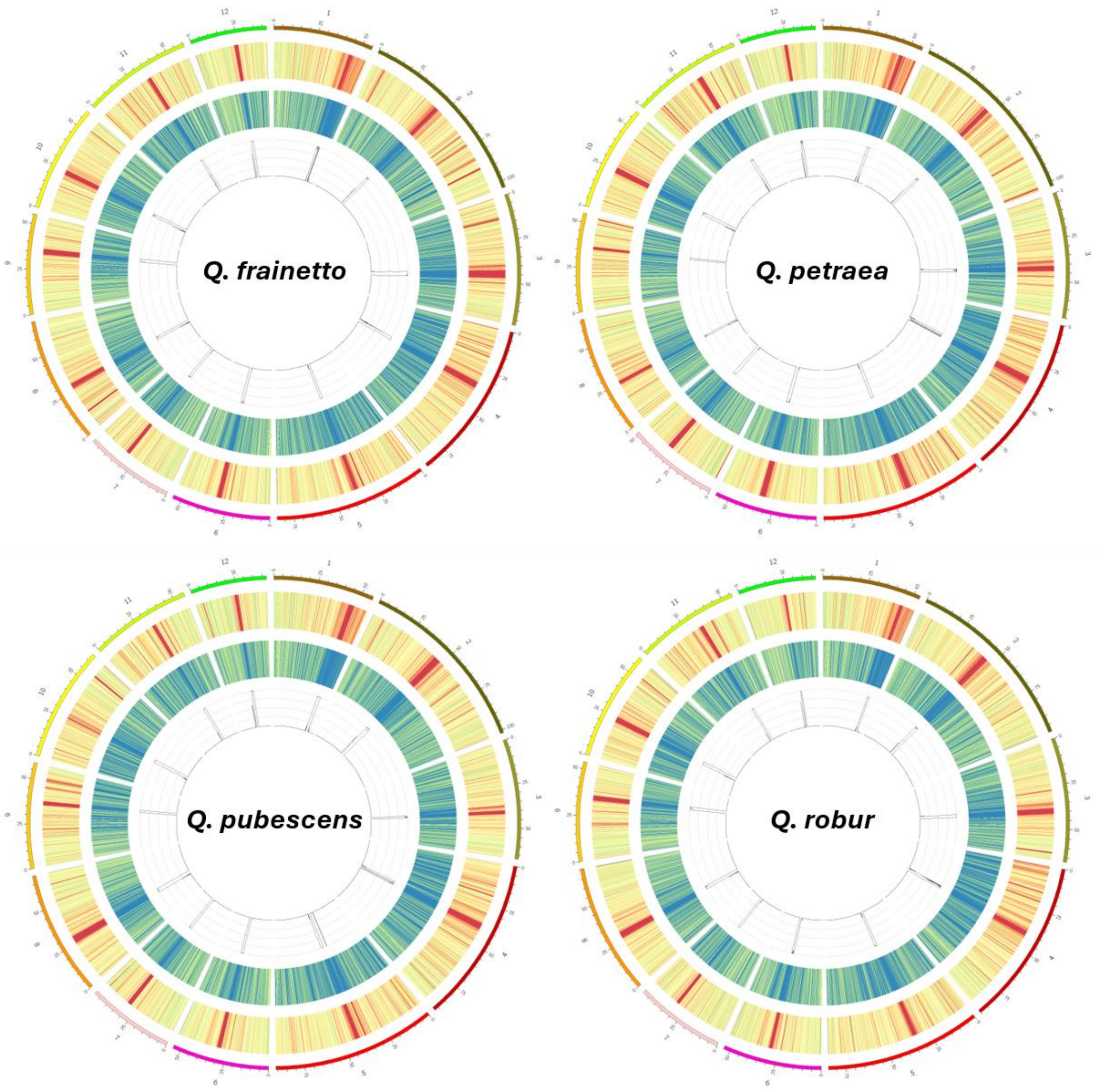


**Supplementary Figure 2. Chromosome-wide distribution of repetitive, coding, and centromeric sequences in haplotype 2 assemblies of *Quercus*.** Circos plots showing the genomic organization of *Q. frainetto*, *Q. petraea*, *Q. pubescens* and *Q. robur* (top-left to bottom-right). Chromosomes are represented by the outer colored ideograms and numbered according to the reference genome assembly. The outer heatmap represents repeat density, with the color scale ranging from yellow (lower density) to red (higher density), whereas the inner heatmap represents gene density, with the color scale ranging from blue (lower density) to yellow (higher density). The innermost histogram represents the abundance of centromeric repeat sequences. All features were calculated in non-overlapping 100 Kb genomic windows.

**Supplementary Table 1. RNA sequencing datasets used in this study.**

| accession | bioproject | biosample | instrument model | library strategy | species | tissue |
|---|---|---|---|---|---|---|
| SRR33253989 | PRJNA1254183 | SAMN48100713 | Illumina NovaSeq 6000 | RNA-Seq | *Quercus robur* | Taproot, control condition |
| SRR33253984 | PRJNA1254183 | SAMN48100708 | Illumina NovaSeq 6000 | RNA-Seq | *Quercus robur* | Taproot, drought stress |
| SRR33253987 | PRJNA1254183 | SAMN48100711 | Illumina NovaSeq 6000 | RNA-Seq | *Quercus robur* | Lateral roots, control condition |
| SRR33253982 | PRJNA1254183 | SAMN48100706 | Illumina NovaSeq 6000 | RNA-Seq | *Quercus robur* | Lateral roots, drought stress |
| SRR30675826 | PRJNA1161636 | SAMN43784840 | Illumina NovaSeq 6000 | RNA-Seq | *Quercus robur* | leaves |
| SRR30675825 | PRJNA1161636 | SAMN43784839 | Illumina NovaSeq 6000 | RNA-Seq | *Quercus robur* | leaves |
| SRR26195204 | PRJNA1021581 | SAMN37565332 | Illumina NovaSeq 6000 | RNA-Seq | *Quercus robur* | mature leaf |
| SRR12717471 | PRJNA665779 | SAMN16268537 | Illumina HiSeq 3000 | RNA-Seq | *Quercus robur* | white roots |
| ERR3348510 | PRJEB32849 | SAMEA5682743 | Illumina HiSeq 2000 | RNA-Seq | *Quercus robur* | Developing bud |
| ERR3348518 | PRJEB32849 | SAMEA5682751 | Illumina HiSeq 2000 | RNA-Seq | *Quercus robur* | Developing bud |
| ERR359852 | PRJEB4873 | SAMEA2229347 | Illumina HiSeq 2000 | RNA-Seq | *Quercus robur* | quiescent buds |
| ERR359855 | PRJEB4873 | SAMEA2229347 | Illumina HiSeq 2000 | RNA-Seq | *Quercus robur* | callus |
| ERR359850 | PRJEB4873 | SAMEA2229347 | Illumina HiSeq 2000 | RNA-Seq | *Quercus robur* | xylem |
| SRR9937109 | PRJNA381515 | SAMN12541762 | NextSeq 500 | RNA-Seq | *Quercus pubescens* | leaf spring |
| SRR9937082 | PRJNA381515 | SAMN12541780 | NextSeq 500 | RNA-Seq | *Quercus pubescens* | leaf summer |
| SRR22815330 | PRJNA910851 | SAMN32236625 | Illumina HiSeq 4000 | RNA-Seq | *Quercus petraea* | fresh young leaves or buds |
| ERR6720554 | PRJEB19536 | SAMEA10058237 | Illumina HiSeq 2000 | RNA-Seq | *Quercus petraea* | white roots, control sample |
| ERR1981092 | PRJEB17876 | SAMEA4545696 | Illumina HiSeq 2000 | RNA-Seq | *Quercus petraea* | buds |
| ERR1854534 | PRJEB17876 | SAMEA4545696 | Illumina HiSeq 2000 | RNA-Seq | *Quercus petraea* | ecodormant buds |
| SRR22815341 | PRJNA910851 | SAMN32236615 | Illumina HiSeq 4000 | RNA-Seq | *Quercus frainetto* | fresh young leaves or buds |
